# The TARZN complex binds *de novo* enhancer mutations and promotes oncogenic expression in T-ALL

**DOI:** 10.1101/2025.01.24.634708

**Authors:** Nurkaiyisah Zaal Anuar, Chai Yeen Goh, Boon Haow Chua, Shi Hao Tan, Joana R. Costa, Marc R. Mansour, Takaomi Sanda, Dennis Kappei

**Author notes:** Correspondence should be addressed to Dennis Kappei. These authors contributed equally (Nurkaiyisah Zaal Anuar, Chai Yeen Goh).

## Abstract

TAL1 is overexpressed in 40-60% of T-cell acute lymphoblastic leukemia (T-ALL) cases and forms an oncogenic core regulatory circuit (CRC) with other transcription factors such as LMO1, LMO2 and GATA3. In 5% of T-ALL cases an insertion of a consensus GT dinucleotide (MuTE) is observed upstream of the *TAL1* gene, driving TAL1 overexpression. Using an *in vitro* reconstitution DNA pull-down assay combined with quantitative mass spectrometry, we identified proteins that preferentially bound to the MuTE sequence and demonstrated that among the candidates the RNA methyltransferase TARBP1 and the zinc finger proteins ZBTB2, ZBTB25 and ZNF639 form a complex that we term TARZN. Interestingly, the TARZN complex also bound to *de novo* super enhancer sites upstream of the *LMO1* and *LMO2* genes in T-ALL cells, indicating a putative common mechanism between these different non-coding driver mutations. Furthermore, knock-down of all TARZN members resulted in lower TAL1 protein expression in MuTE-positive but not in MuTE-negative T-ALL cells. Given TARZN’s methyltransferase activity and the lack of concomitant *TAL1* mRNA level changes, we investigated reduced TAL1 translation and identified reduced neo-synthesised TAL1 protein levels upon TARBP1 knockdown. Overall, these data suggest that the TARZN complex promotes oncogenic expression in T-ALL via co-transcriptional RNA methylation.

## INTRODUCTION

T-cell acute lymphoblastic leukemia (T-ALL) is an aggressive hematological cancer characterised by an excessive production of immature T-cell progenitors in the thymus^1,2^. TAL1, a basic Helix-Loop-Helix (bHLH) transcription factor, is one of the major oncogenic drivers of T-ALL and is aberrantly upregulated in 40-60% of T-ALL cases^3,4^. TAL1 along with its regulatory partners HEB, E2A, LMO1, LMO2, GATA3, and RUNX1 form a complex that regulates their own transcription through a positive feedback loop, as well as the expression of multiple downstream targets, of which many are genes involved in T-cell development and growth^5^. The overexpression of TAL1 results in a differentiation block that leads to the malignant transformation of precursor T-cells^5,6^ and is therefore critical for T-ALL cell maintenance. Indeed depletion of TAL1 and/or its direct downstream targets have been shown to significantly retard T-ALL cell growth^7–10^.

Several genetic alterations that lead to TAL1 overexpression in T-ALL have been reported, with the most well-known being a ∼90 kb deletion upstream of *TAL1* reported in ∼30% of T-ALL cases, which puts *TAL1* under the regulatory element of *SIL*, an ubiquitously expressed gene located upstream of *TAL1*^11,12^. In ∼10% of T-ALL cases, chromosomal translocation of the *T-cell Receptor (TCR)* element results in fusion between *TCR* and *TAL1* genes, leading to hijacking of the *TCR* regulatory element to drive *TAL1* expression^4,13,14^. Finally, recurrent somatic mutations within the non-coding region ∼7.5 kb upstream of *TAL1* found in ∼5% of T-ALL cases were reported to drive oncogenic TAL1 expression^15^. These mutations comprise small insertions of a few nucleotides upstream of the *TAL1* gene, which form the consensus of a GT dinucleotide insertion termed as “<u>Mu</u>tation of the <u>T</u>AL1 <u>E</u>nhancer” (MuTE). The insertion introduces a MYB binding motif and leads to the formation of a widespread H3K27ac super enhancer signature around the MuTE site. MYB binds to the *de novo* motif alongside CBP and the TAL1 transcriptional complex to initiate the aberrant transcriptional activity that leads to TAL1 overexpression^15^. In agreement with this, knockdown of MYB resulted in lower TAL1 expression. However, considering the nearly complete loss in TAL1 expression upon the excision of the MuTE allele^15^ along with the nature of super enhancers to recruit many transcription factors^16^, we reasoned that there are likely other proteins to contribute to MuTE-mediated TAL1 expression. In addition to MuTE, several other *de novo* mutations in the non-coding regions associated with *LMO1* and *LMO2*, two other T-ALL oncogenes, were subsequently reported^17–19^. Like MuTE, these mutations also introduce MYB binding motifs and lead to super enhancer formation that drives the overexpression of these oncogenes^17–19^, which raises the possibility that similar additional regulatory factors may also be in play at these mutant sites.

In this study, we report the binding of the TARZN complex to the MuTE, LMO1 and LMO2 *de novo* super enhancer sites. Loss-of-function experiments demonstrated that the TARZN complex is involved in their oncogenic expression regulation in mutation-positive T-ALL cells.

## RESULTS

### Quantitative mass spectrometry identified TAL1 MuTE-specific proteins

To systematically identify MuTE-specific proteins, we performed *in vitro* reconstitution DNA pull-downs by incubating biotinylated oligonucleotides containing the WT or the MuTE consensus sequence (GT insertion) with MOLT3 nuclear protein extracts harbouring this exact mutation. Quadruplicates of each pull-down were analysed using label-free quantitative (LFQ) mass spectrometry (MS)^20^, and proteins with MuTE-to-WT enrichment ratios of 4-fold or above and p-values < 0.01 were considered as MuTE-specific binders (**Fig. 1A**). In total, we identified 11 proteins specific to the MuTE sequence (**Fig. 1B, Supplementary Table 1**). To eliminate possible cell line-specific effects, we performed the same experiment using nuclear protein extracts from Jurkat cells that carry a 12-bp insertion containing the consensus GT insertion^15^ and identified nine overlapping proteins (**Fig. 1C, Supplementary Table 2**). To further substantiate these results, we repeated pull-downs with both Jurkat and MOLT-3 extracts using <u>S</u>table Isotope <u>L</u>abelling with <u>A</u>mino acids in <u>C</u>ell culture (SILAC)^21^ as an independent quantitation method for the mass spectrometry analysis (**Fig. S1A**). Here, only proteins with both ‘forward’ and ‘reverse’ SILAC ratios of 4 and above are considered as specific binders. Once again, a similar list of proteins to those identified by LFQ was captured in both SILAC datasets (**Fig. S1B-D, Supplementary Tables 3 and 4**). Overall, we identified ten proteins that consistently (in at least three out of four experiments) bound to the MuTE over the WT sequence across both cell lines (**Fig. S1D**), including several zinc finger proteins (ZBTB2, ZBTB25, ZNF639, ZNF652), Jerky protein homolog-like (JRKL) and its paralogous interactor Tigger transposable element-derived protein 2 (TIGD2), and Lethal(3)malignant brain tumor-like protein 3 (L3MBTL3), all with predicted DNA binding ability^22,23^. Our candidates also included Homeobox-containing protein 1 (HMBOX1), known to bind to the telomeric repeat sequence TTAGGG^24^ along with Atherin (SAMD1) that is an interactor of L3MBTL3^25^, and the RNA methyltransferase TAR (HIV-1) RNA binding protein 1 (TARBP1)^26,27^. Of note, while MYB was previously described to be present at the MuTE locus *in vivo*^15^, we had not detected MYB in any of our MS experiments (**Fig. 1C**).

**Figure 1.**
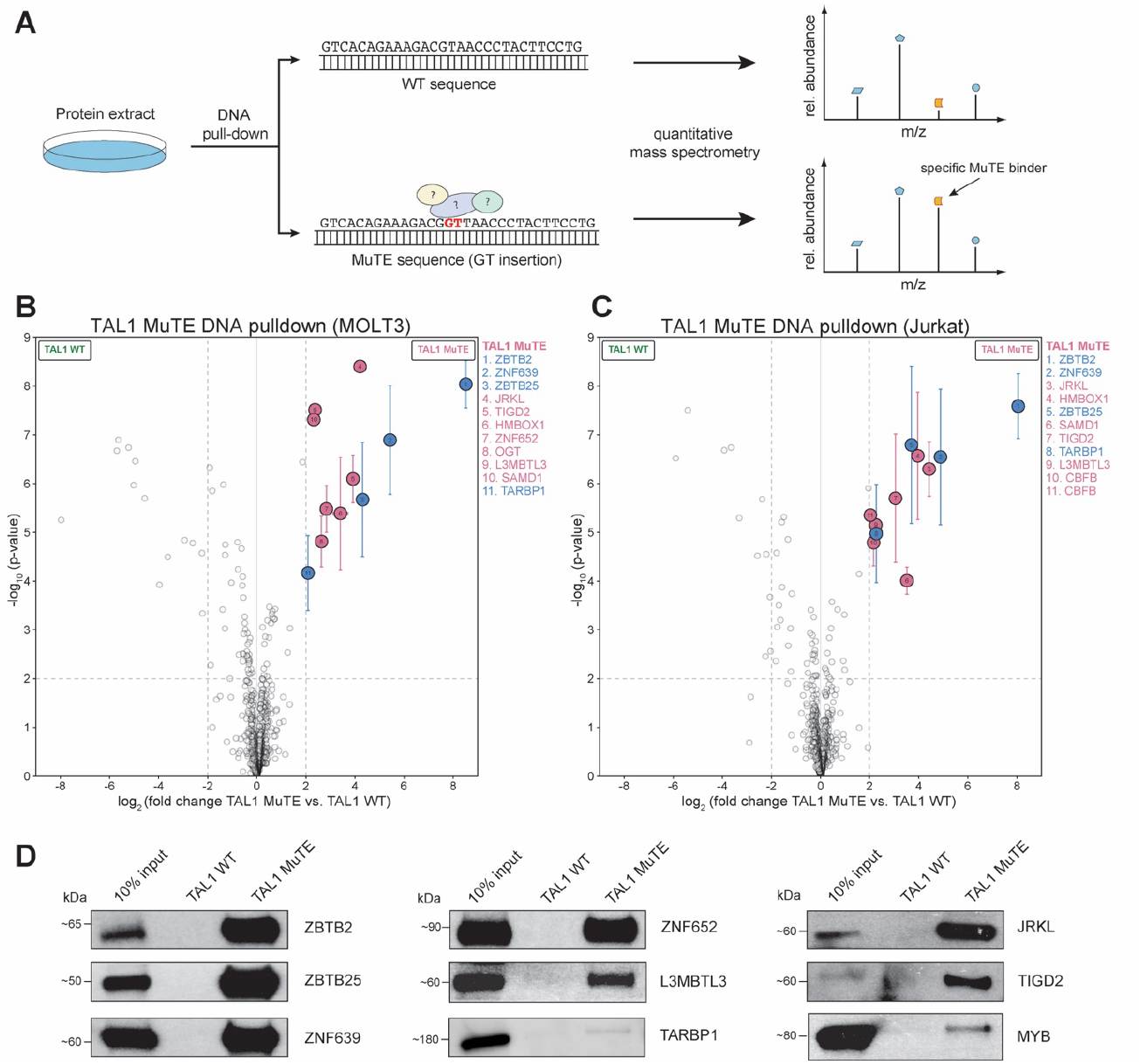
Identification of TAL1 MuTE-specific proteins through quantitative MS. (A) Label-free nuclear protein extracts are reconstituted with either WT or MuTE oligonucleotides *in vitro*. Specific interaction partners are differentiated from background binders by a ratio significantly different from 1:1. All pull-downs were performed in four replicates and analysed by quantitative label-free proteomics. (B) Volcano plot of DNA pull-down with nuclear protein extracts from MOLT-3 and (C) Jurkat cells. Specifically enriched proteins (numbered circles) are distinguished from background binders by a two-dimensional cut-off of >4-fold enrichment and p<0.01. Two-dimensional error bars represent the standard deviation based on iterative imputation cycles during the label-free analysis to substitute zero values (e.g. no detection in the WT pull-downs). TARZN complex members are colored in blue. (D) Immunoblots confirming specific binding of candidate proteins and MYB to the TAL1 MuTE sequence in comparison to the WT sequence using nuclear protein extracts from Jurkat cells. Blots shown are representative results from at least three independent experiments.

We were able to validate specific binding to the MuTE sequence in our DNA pull-down assays for eight of our candidate proteins via immunoblotting (**Fig. 1D**), while the remaining two (HMBOX1 and SAMD1) could not be corroborated due to the lack of a functional commercial antibody. All eight proteins showed specific binding to the MuTE sequence in comparison to the WT sequence as captured in our MS analysis. Most candidates showed strong absolute enrichment when compared to the band intensity of their 10% input signal, with the exception of L3MBTL3 and TARBP1 which is consistent with their low coverage in our mass spectrometry experiments (**Fig. 1D, Fig. 1B-C, S1B-C, Supplementary Tables 1-4**). To differentiate between a lack of enrichment in the DNA pull-down vs. simple non-identification in our MS analyses for technical reasons, we included MYB in our immunoblot analysis. Here, MYB was indeed preferentially bound to the MuTE sequence but with significantly weaker absolute enrichment than other identified proteins, comparable to the weak enrichment of L3MBTL3 and TARBP1 (**Fig. 1D**). This combination of specific binding with weak absolute enrichment could potentially explain the lack of MYB identification in our MS analyses.

### ZBTB2, ZBTB25, ZNF639 & TARBP1 bind to multiple mutation-dependent *de novo* super enhancer mutations

The expression of TAL1 and its downstream targets in hematopoietic cells is largely regulated by the TAL1 complex consisting of TAL1, HEB, E2A, LMO1/2, GATA3, and RUNX1^5,28^. Notably, *TAL1, LMO1*, and *LMO2* loci were found to harbour multiple non-coding mutations. Several of these mutations also introduce *de novo* MYB binding motifs and promote super enhancer formation that in return drives the overexpression of these oncogenes^8,17–19,29^. To identify potential common factors between these mutant sequences, we applied our DNA pull-down strategy coupled with quantitative MS analysis to a C-to-T point mutation upstream of the *LMO1* gene in Jurkat cells (**Fig. 2A**) using SILAC-labelled Jurkat nuclear protein extracts (**Fig. S1A**). Interestingly, some of the MuTE-specific proteins were also enriched on the mutant T-allele sequence, including ZBTB2, ZBTB25, ZNF639 and TARBP1 (**Fig. 2B, Supplementary Table 5**). Similarly to the MuTE sequence, we validated specific binding to the LMO1 mutant sequence in DNA pulldown assays for these candidates via immunoblotting, where ZBTB2, ZBTB25, ZNF639 and TARBP1 showed preferential binding to the C-to-T mutant sequences over the WT sequences with a similar enrichment profile (**Fig. 2C**).

**Figure 2.**
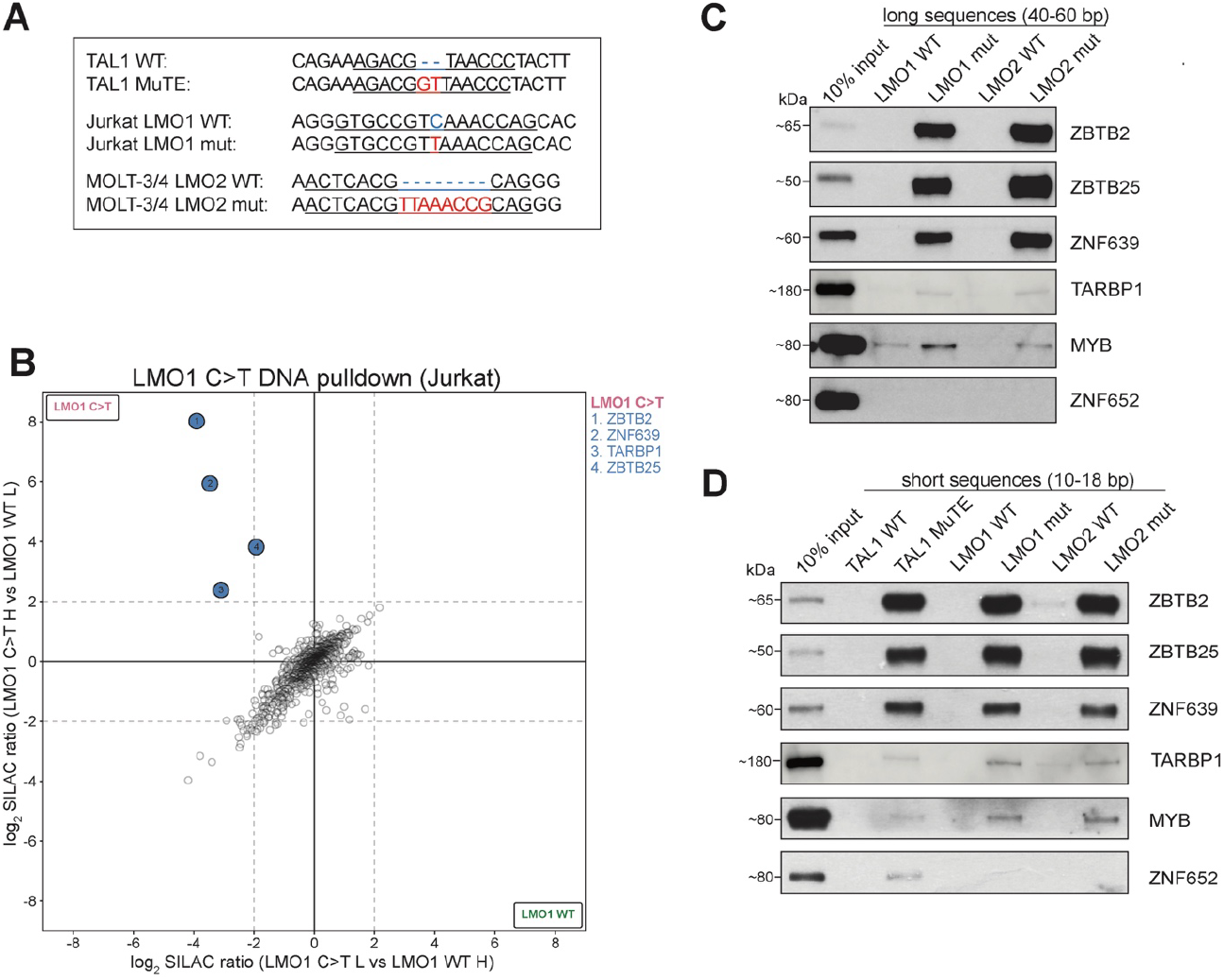
ZBTB2, ZBTB25, ZNF639 and TARBP1 bind to multiple enhancer mutations. (A) Mutation-dependent super enhancer sites upstream of TAL1, LMO1, and LMO2 containing *de novo* MYB binding sites. WT sequences are highlighted in blue and mutations in red. (B) Scatterplot of DNA pull-downs with SILAC-labelled nuclear protein extracts from Jurkat cells. Heavy-labelled SILAC extracts were applied to the C-to-T mutant oligonucleotides and light-labelled extracts to the WT oligonucleotides (y-axis) or vice versa (x-axis) and log_2_ SILAC ratios are displayed. Specific mutant-binders are found in the upper left quadrant as numbered (cut-off ≥4-fold enrichment). TARZN complex members are colored in blue. (C) Immunoblots showing specific binding of the trimeric zinc finger complex to super enhancer-inducing mutant sequences with oligonucleotides containing the long and (D) short sequences using Jurkat nuclear protein extracts. MYB and ZNF652 were included for comparison. ZNF652 served as a control that binds only to the MuTE but not to the other sequences. Blots shown are representative results from at least three independent experiments. H: heavy, L: light, WT: wild-type, mut: mutant.

We also compared binding of ZBTB2, ZBTB25, ZNF639 and TARBP1 to another super enhancer-inducing mutation consisting of a 8-bp insertion upstream of the *LMO2* gene in MOLT-3/4 cells^17^ (**Fig. 2A**). As with the MuTE and LMO1 mutant sequences, ZBTB2, ZBTB25, ZNF639 and TARBP1 preferentially bound to the *LMO2*-associated mutant 8-bp insertion sequence compared to the WT control (**Fig. 2C**). Again, MYB only displayed weak absolute enrichment to all mutant sequences.

Our DNA pull-down oligonucleotide probes thus far consisted of 40-60 bp including flanking sequences around the mutation sites (the ‘long’ sequences, **Supplementary Table 6**). To pinpoint whether the binding is confined to the respective mutations or requires the flanking sequences, we generated oligonucleotides containing a maximum of 7 bp up- and downstream of the respective mutation (underlined in **Fig. 2A**, ‘short sequences’ in **Supplementary Table 6**). Consistent with the previous pull-down experiments, ZBTB2, ZBTB25, ZNF639 were again strongly enriched on all three super enhancer-inducing motifs as compared to their respective WT sequences (Fig. 2D). Also similar to our previous data, TARBP1 and MYB showed preferential binding to all three short mutant sequences, albeit with weak absolute signal. Overall, the weak absolute enrichment of TARBP1 at these mutant sites suggests that TARBP1 may act as indirect binder, due to lack of a DNA binding domain. In sum, we identified that ZBTB2, ZBTB25, ZNF639 and TARBP1 recurrently bind to super enhancer-forming mutant sequences upstream of critical T-ALL oncogenes.

### ZBTB2, ZBTB25, ZNF639 & TARBP1 form the TARZN complex

Previously, ZBTB2 was identified as a direct binder and regulator of CpG island promoters involved in mouse embryonic stem cell differentiation and was shown to form a complex with ZBTB25 and ZNF639^30^. Given this prior knowledge and that ZBTB2, ZBTB25, ZNF639 and TARBP1 recurrently bound both the MuTE as well as the LMO1 and LMO2 mutations, we next asked whether these four proteins interact with each other in T-ALL cells. To test this, we analysed immunoprecipitations (IP) by quantitative mass spectrometry in Jurkat cells overexpressing either ZBTB2, ZBTB25, ZNF639 or TARBP1 with a C-terminal FLAG sequence (**Fig 3A**). In all reciprocal IP-MS experiments, we were able to significantly enrich all four proteins (**Fig. 3B-D, Supplementary Tables 7-10**), establishing ZBTB2, ZBTB25, ZNF639 and TARBP1 as a tetrameric complex that we term TARZN (TARBP1-zinc finger complex). This was further validated by immunoblotting (**Fig. S2A**), which showed strong mutual enrichment among the zinc fingers but relatively weaker enrichment of TARBP1. These IP-MS results (**Fig. S2B**) combined with TARBP1’s weak absolute enrichment in the DNA pulldown assays and the lack of a DNA-binding domain (**Fig. 1D, 2C-D**) suggest that the three zinc fingers may act to recruit TARBP1.

**Figure 3.**
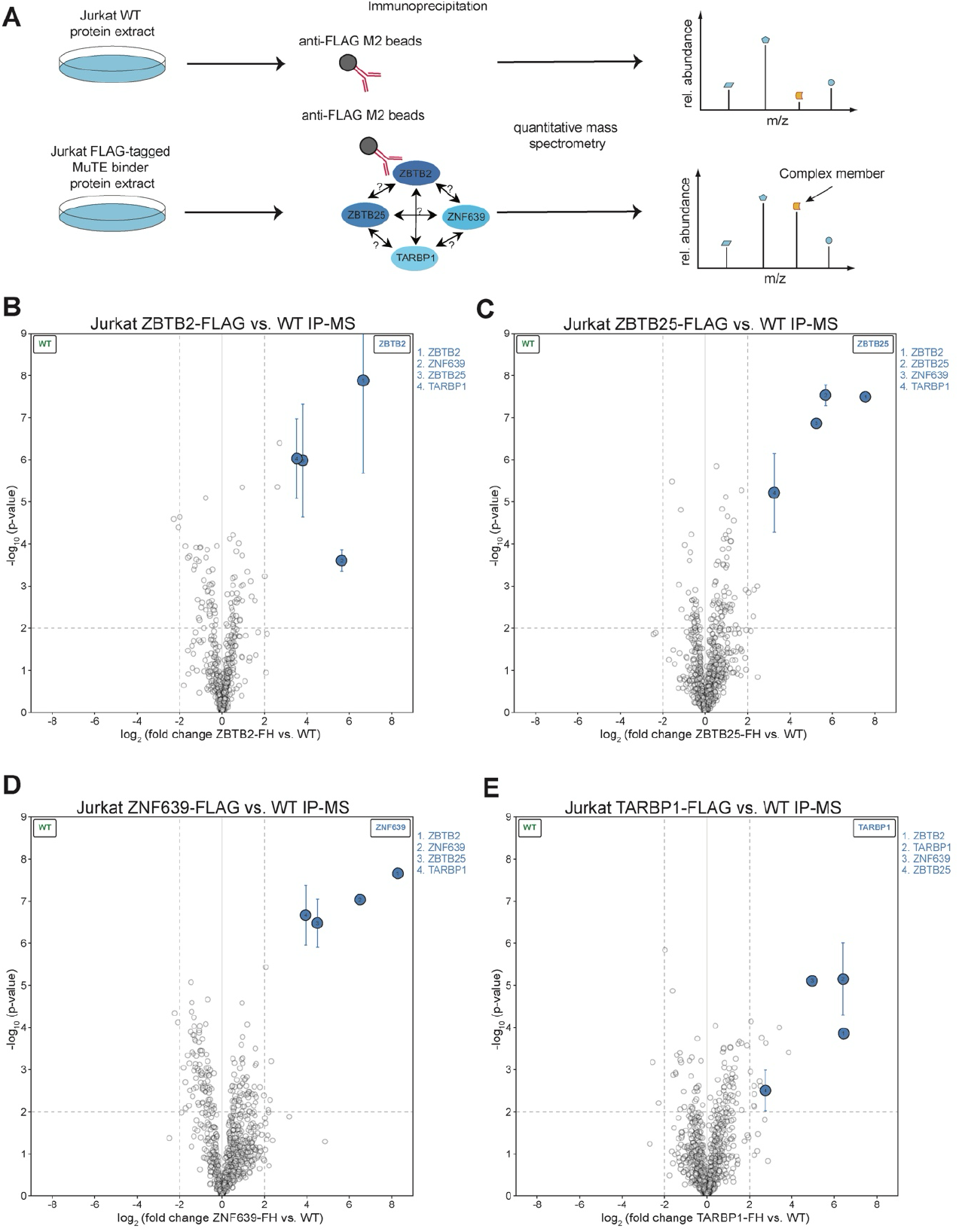
ZBTB2, ZBTB25, ZNF639 & TARBP1 form the TARZN complex. (A) Nuclear protein extracts from either Jurkat WT or Jurkat cells overexpressing FLAG-tagged ZBTB2, ZBTB25, ZNF639 or TARBP1 were incubated with Anti-FLAG M2 magnetic beads. Specific interaction partners are distinguished from background binders by a two-dimensional cut-off of >4-fold enrichment and p<0.01. All pull-downs were performed in four replicates and analysed by quantitative label-free proteomics. Volcano plots comparing immunoprecipitation from Jurkat WT vs ZBTB2-FLAG (B), ZBTB25-FLAG (C), Jurkat ZNF639-FLAG (D) or TARBP1-FLAG (E).

### The TARZN complex regulates TAL1 protein expression

To test whether MuTE-specific binding of the TARZN complex translates to a functional impact on TAL1 expression in T-ALL cells, we knocked down *ZBTB2, ZBTB25, ZNF639* and *TARBP1* mRNAs in Jurkat cells using shRNAs. While little to no effect on *TAL1* transcript levels was observed upon knockdown of the zinc fingers, knockdown of TARBP1 exhibited a 40-50% reduction in *TAL1* mRNA levels, similar to *MYB* knockdown (**Fig. 4A, S3A**). In contrast, knockdowns of all members of the TARZN complex led to a clear reduction in TAL1 protein levels (**Fig. 4B**) similar to the effect of MYB depletion (**Fig. S3B**). To confirm that this effect was mediated by the MuTE locus, we performed the same knockdown experiment in CCRF-CEM cells, a TAL1-positive MuTE-negative T-ALL cell line whose TAL1 overexpression is driven by the ∼90 kb *SIL-TAL* deletion^5,11,31^. Indeed knocking down all members of the TARZN complex did not affect TAL1 protein expression in these cells (**Fig. 4C**) in agreement with a MuTE-specific regulation. In contrast, MYB knockdown led to a significant decrease in TAL1 protein levels in CCRF-CEM (**Fig. S3C**). Furthermore, to validate the role of the TARZN complex at other *de novo* super enhancer sites, we performed knockdown of ZBTB25 in MOLT-3 cells (which contain the LMO2 *de novo* enhancer mutation). Similar to the MuTE-dependent effect on TAL1 expression, we observed little to no effect on LMO2 transcript levels (**Fig. S3D**) but a significant attenuation of LMO2 protein levels (**Fig. S3E**). Overall, these data suggest that the TARZN complex promotes an oncogenic gene expression program in T-ALL based on recurrent *de novo* enhancer mutations.

**Figure 4.**
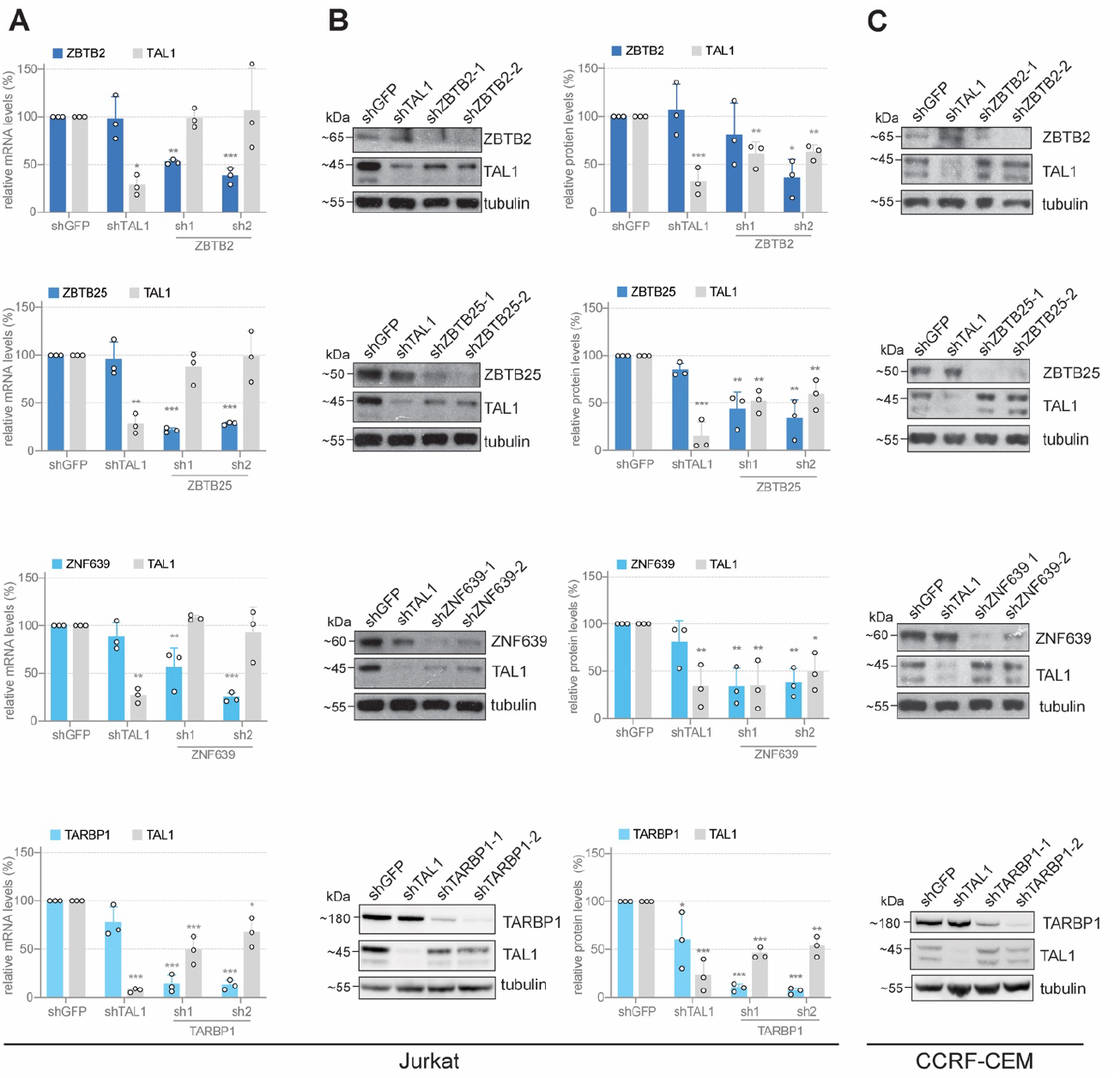
The TARZN complex regulates TAL1 protein expression. (A) qPCR data measuring mRNA knockdown of MuTE binders and their effect on *TAL1* expression at the mRNA level in Jurkat cells. Cells were transduced with lentivirus expressing shRNAs of the target proteins, and puromycin selection (0.75 µg/ml) was initiated 48 h post-transduction for a further 72 h. (B) Immunoblots showing shRNA-mediated knockdown of *ZBTB2, ZBTB25, ZNF639* and *TARBP1* in Jurkat cells and their effect on TAL1 protein expression. Whole cell lysates were used as the input for immunoblotting. Right panel shows quantification of the immunoblots from Jurkat cells performed using ImageJ. (C) Immunoblots showing shRNA-mediated knockdown of ZBTB2, ZBTB25, ZNF639 and TARBP1 in MuTE-negative CCRF-CEM cells and their effect on TAL1 protein expression. Experiment was performed as described for Jurkat cells in (B). All data shown depicts the mean±SD of three biological replicates. Asterisks represent statistical significance (^*^ p≤0.05, ^**^ p≤0.01, ^***^ p≤0.001) calculated by one-way ANOVA followed by Dunnett’s multiple comparison tests.

### TARBP1 regulates translation efficiency of TAL1

Given that in TARBP1 the TARZN complex contains a RNA 2’-O-methyltransferase and the effect on TAL1 expression was largely confined to pronounced changes in TAL1 protein levels, we hypothesised that the TARZN complex may methylate TAL1 mRNA and impact protein translation, in line with previous work implicating 2’-O-methylation (Nm) in translation regulation^32^. To test this, we performed bio-orthogonal non-canonical amino acid tagging (BONCAT) using L-azidohomoalanine (AHA)^33^, an analog of L-methionine, allowing us to specifically label neo-synthesised proteins. This AHA tag is then coupled to desthiobiotin by click chemistry followed by capture with streptavidin beads to specifically quantify changes in neo-synthesised TAL1 protein levels following TARBP1 knockdown (**Fig. 5A**). Indeed, neo-synthesised TAL1 protein was only detected in AHA-treated samples but not DMSO-treated controls (**Fig. 5B**) and TARBP1 knockdown resulted in a significant reduction of neo-synthesised TAL1 levels (**Fig. 5B-C**). This reduction was quantitatively similar to the loss of TAL1 protein in steady-state experiments (**Fig. 4B**), suggesting that reduced TAL1 translation indeed explains the loss of TAL1 protein levels upon depletion of TARZN complex members. Based on these results, we propose a model in which the TARZN complex is recruited to the *TAL1* locus in a MuTE-dependent manner and promotes TAL1 translation via co-transcriptional methylation of *TAL1* mRNA (**Fig. 5D**).

**Figure 5.**
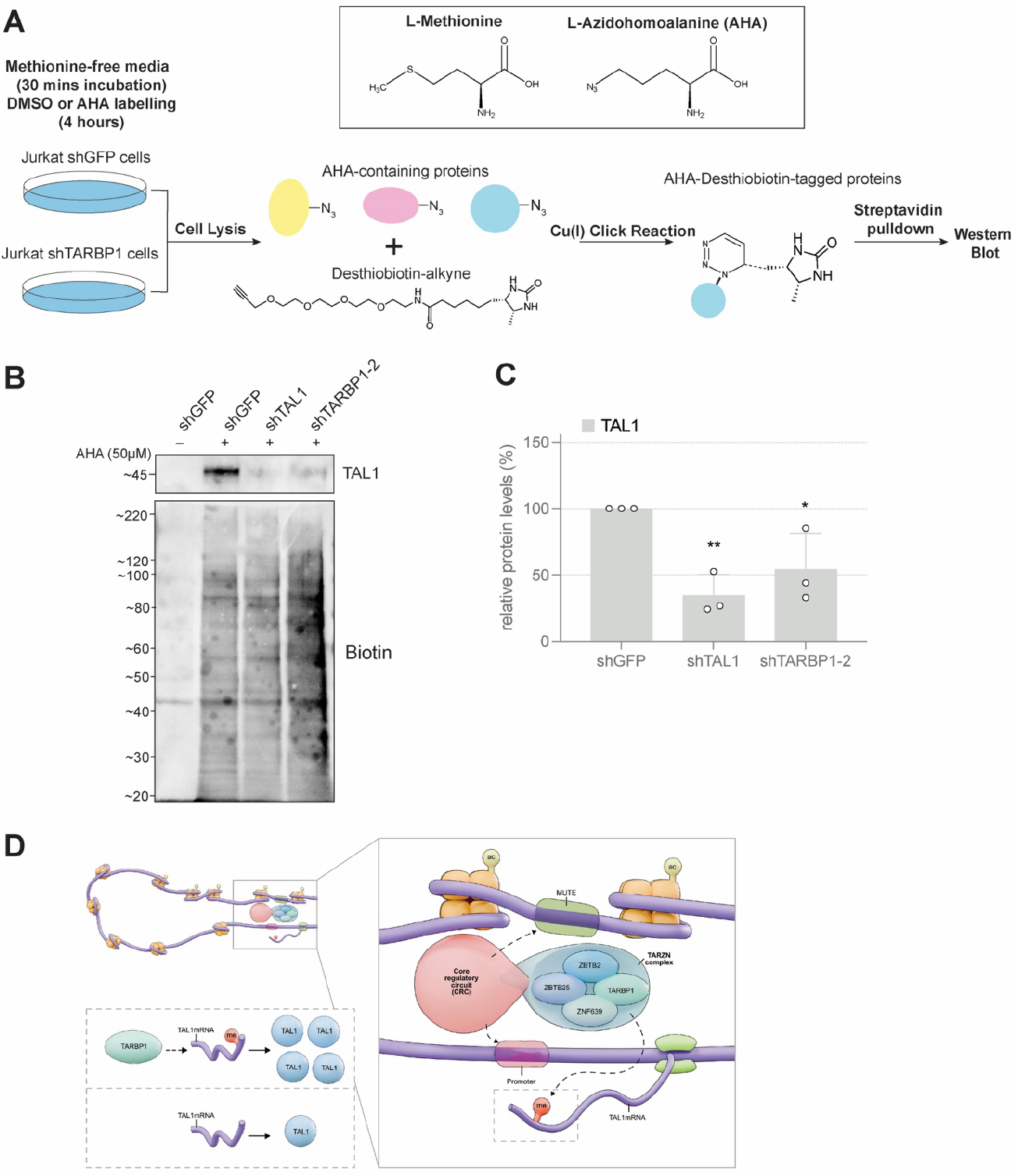
TARBP1 regulates translation efficiency of TAL1. (A) Schematic of the BONCAT workflow including AHA labelling and biotin tagging prior to streptavidin pull-down to enrich neo-synthesised proteins. (B) AHA-deshthiobiotin pull-down upon shRNA-mediated knockdown of TARBP1 in Jurkat cells and its effect on neo-synthesised TAL1 protein levels (top) with validation of the biotin labelling efficiency (bottom). Cells were transduced with lentivirus expressing shRNAs of the target proteins, and puromycin selection (0.75 µg/ml) was initiated 48 h post-transduction for a further 72 h. (C) Quantification of the immunblot in Fig. 5B depicting mean±SD of three biological replicates. Asterisks represent statistical significance (^*^ p≤0.05, ^**^ p≤0.01) calculated by one-way ANOVA followed by Dunnett’s multiple comparison tests. (D) Working model of co-transcriptional TAL1 mRNA methylation and protein expression regulation via the TARZN complex in MuTE-positive T-ALL cells.

## DISCUSSON

TAL1 overexpression has long been recognised as one of the hallmarks in T-ALL^1,4^. The MuTE insertion is one mechanism to drive oncogenic TAL1 expression by creating a super enhancer upstream of the TAL1 gene, with TAL1 and their core regulatory circuit (CRC) members recruited to the MuTE site *in vivo*^15^. However, the identification of these factors was based on sequence motifs and binding prediction, which by default requires the binding factors to be well-characterised. Such predictive *in silico* methods are likely to ignore putative additional factors that have been less well characterised and for which the necessary ChIP-seq data is unavailable, which is indeed the case for the majority of proteins identified in our proteomics screen. In addition, parallel work analysing MYB-containing chromatin complexes by MYB ChIP-MS suggests that the MYB antibody used to validate binding to the MuTE locus may have recognised additional proteins, questioning whether MYB is indeed a physical member of the TAL1 CRC^34^. This may explain the comparably weak absolute enrichment of MYB in our DNA pull-down assays and point to a scenario in which other TAL1 CRC members directly bind to sequence elements adjacent to the MuTE, LMO1 and LMO2 mutations.

Indeed, in this study we captured multiple novel MuTE-specific proteins by combining an *in vitro* reconstitution DNA pull-down assay with quantitative MS. Apart from the MuTE insertion, several other mutations have been described that also introduce *de novo* MYB binding motifs and form super enhancers that drive the overexpression of other critical T-ALL oncogenes, *LMO1* and *LMO2*, in T-ALL cells^17,18^. The LMO1 and LMO2 proteins do not directly bind DNA but serve as important mediator proteins that hold the TAL1 regulatory complex together^18^. Intriguingly, ZBTB2, ZBTB25, ZNF639 and TARBP1 were found to bind to all three *de novo* super enhancer sequences that we tested. Consistent across the different non-coding driver mutations, the zinc fingers demonstrated strong specific enrichment at the mutant sites (**Fig. 1D, 2C**) as well as a strong association with each other (**Fig. S2B**), which is in agreement with prior work establishing ZBTB2, ZBTB25, and ZNF639 as a trimeric zinc finger complex involved in the regulation of mouse embryonic stem cell differentiation^30^. In contrast, TARBP1 consistently exhibited weaker enrichment at the mutant sites (**Fig. 1D, 2C**) and overall a weaker association with the zinc fingers (**Fig. S2B**). This presents a scenario wherein the three zinc fingers bind directly to the DNA in a sequence-specific manner and recruit TARBP1 to enable locus-specific methyltransferase activity in analogy to how transcription factors are frequently the sequence-specific adaptor proteins that recruit epigenetic complexes and hence their individual enzymatic activities.

TARBP1 has been predicted to be part of the Spou-TrmD (SPOUT) methyltransferase family of RNA methyltransferases, which methylates the 2’-hydroxyl on the ribose moiety, otherwise known as 2’-O-methylation (Nm)^27^. The Nm modification has an abundance of 0.5% per mRNA transcript^32^, and has been shown to regulate changes in mRNA stability and translation efficiency^32,35^. In our study we demonstrated changes in TAL1 protein levels upon knockdown of the TARZN complex but little to no change in mRNA levels, which we can attribute to an impeded translation efficiency. We further corroborated these findings by profiling changes in neo-synthesised TAL1 levels, which demonstrated a significant reduction in neo-synthesised TAL1 levels following TARBP1 knockdown. Overall, our findings support a model, in which ZBTB2, ZBTB25 and ZNF639 recruit TARBP1 to the MuTE locus. TARBP1 then methylates TAL1 mRNA co-transcriptionally, thus affecting the TAL1 translation efficiency, without prominently affecting mRNA stability. These data represent a second parallel mechanism contributing to overall TAL1 levels in addition to the well-established transcriptional regulation via the TAL1 CRC components and may potentially provide a new targetable vulnerability in T-ALL.

## METHODS

### Cell culture

Jurkat, MOLT-3, and CCRF-CEM cells were cultured in RPMI 1640 (Gibco) medium supplemented with 10% fetal bovine serum (Gibco) and 100 U/mL penicillin-streptomycin (Gibco). HEK293T cells were cultured in Dulbecco’s Modified Eagle’s Medium (DMEM) containing 4.5 g/l glucose, 4 mM glutamine, 1 mM sodium pyruvate and supplemented with 10% fetal bovine serum (Gibco) and 100 U/mL penicillin-streptomycin (Gibco). For SILAC labelling, Jurkat and MOLT-3 cells were incubated in RPMI (−Arg, -Lys) medium containing 10% dialyzed fetal bovine serum (PAN-Biotech) and 100 U/mL penicillin-streptomycin, supplemented with 84 mg/L ^13^C_6_^15^N_4_ L-arginine and 50 mg/L ^13^C_6_^15^N_2_ L-lysine (Cambridge Isotope) or the corresponding non-labelled amino acids, respectively. Successful SILAC incorporation was verified by in-gel trypsin digestion and MS analysis of ‘heavy’ input samples to ensure an incorporation rate of >98%.

### Nuclear protein extraction

Cells were harvested and nuclear extracts were prepared as previously described^36^. Briefly, cells were pelleted prior to incubation and lysis in hypotonic buffer (10 mM HEPES, pH 7.9, 1.5 mM MgCl2, 10 mM KCl) with a dounce homogenizer (type B pestle), where both Jurkat and MOLT-3 cells were lysed mechanically with 10 strokes. Nuclei were washed in PBS and extracted in hypertonic buffer (420 mM NaCl, 20 mM Hepes, pH 7.9, 20% glycerol, 2 mM MgCl2, 0.2 mM EDTA, 0.1% IGEPAL CA-630 (Sigma), 0.5 mM DTT) for 1h at 4°C on a rotating wheel. Samples were centrifuged at maximum speed for 1h at 4ºC. Protein concentration in the supernatant was quantified using Pierce BCA Protein Assay Kit (Thermo Fisher Scientific) as per manufacturer’s instructions.

### *In vitro* reconstitution DNA pull-down

Biotinylated oligonucleotide probes were generated as previously described^36,37^. Oligonucleotide probes were bound to 250 µg (Western blot) or 750 µg (MS) paramagnetic streptavidin beads (Dynabeads MyOne Streptavidin C1, Thermo Fisher Scientific) before incubation with 100µg (Western blot) or 300-400µg (MS) nuclear protein extracts in PBB buffer (150 mM NaCl, 50 mM Tris-HCl pH 7.5, 5 mM MgCl2, 0.5% IGEPAL CA-630 (Sigma)) in the presence of 20µg (Western blot) or 40µg (MS) sheared salmon sperm DNA (Ambion) for 2 h at 4°C on a rotating wheel. Three PBB buffer washes were performed and bound proteins were eluted in 2x Laemmli buffer and boiled for 5 min at 95°C. For SILAC experiments only, the bead suspension of corresponding samples were combined together in the third wash in a new tube.

### Protein Immunoprecipitation

100µg (Western blot) or 300-400µg (MS) Jurkat nuclear extracts overexpressing ZBTB2, ZBTB25, ZNF639 or TARBP1 with a C-terminal FLAG-tag were incubated with 20 µL (Western blot) or 50 µL (MS) of Anti-FLAG M2 magnetic beads (Sigma-Aldrich) in PBB-Tx buffer (150 mM NaCl, 50 mM Tris-HCl pH 7.4, 1 mM EDTA, 1% Triton-X, 1x cOmplete protease inhibitor (Roche)) for 2 h at 4ºC on a rotating wheel. The beads were washed thrice in PBB-Tx buffer and bound proteins were eluted with 2x Laemmli buffer boiled at 95ºC for 5 min.

### Mass spectrometry data acquisition and analysis

DNA pull-down and IP samples were separated on a 12% NuPAGE Bis-Tris gel (Thermo Fisher Scientific) for 30 min (SILAC) or 10 min (LFQ) at 170 V in 1× MOPS buffer (Thermo Fisher Scientific). The gel was fixed using the Colloidal Blue Staining Kit (Thermo Fisher Scientific). For SILAC samples, each lane was divided into 4 equal fractions, while LFQ samples were not fractionated. For in-gel digestion, samples were destained in destaining buffer (25 mM ammonium bicarbonate; 50% ethanol), reduced in 10 mM DTT for 1h at 56°C followed by alkylation with 55mM iodoacetamide (Sigma) for 45 min in the dark. Tryptic digest was performed in 50 mM ammonium bicarbonate buffer with 2 μg trypsin (Promega) at 37°C overnight. Peptides were desalted on StageTips and analysed by nanoflow liquid chromatography on an EASY-nLC 1200 system coupled to a Q Exactive HF mass spectrometer (Thermo Fisher Scientific). Peptides were separated on a C18-reversed phase PicoFrit column (25 cm long, 75 μm inner diameter; New Objective) packed in-house with ReproSil-Pur C18-AQ 1.9 μm resin (Dr Maisch). The column was mounted on an Easy Flex Nano Source and temperature controlled by a column oven (Sonation) at 40°C. A 105-min gradient from 2 to 40% acetonitrile in 0.5% formic acid at a flow of 225 nl/min was used. Spray voltage was set to 2.2 kV. The Q Exactive HF was operated with a TOP20 MS/MS spectra acquisition method per MS full scan. MS scans were conducted with 60,000 at a maximum injection time of 20 ms and MS/MS scans with 15,000 resolution at a maximum injection time of 50-75 ms. The raw files were processed with MaxQuant^38^ version 1.5.2.8 (DNA pulldown) or MaxQuant^38^ version 2.0.1.0 (IP) with preset standard settings for SILAC labelled including the re-quantify option or LFQ^20^ samples with match-between-runs activated. Carbamidomethylation was set as fixed modification while methionine oxidation and protein N-acetylation were considered as variable modifications. Search results were filtered with a false discovery rate of 0.01. Known contaminants, proteins groups only identified by site, and reverse hits of the MaxQuant results were removed and only proteins were kept that were quantified by SILAC ratios in both ‘forward’ and ‘reverse’ samples, or by LFQ intensity.

### Plasmids and cloning

Full-length ZBTB2, ZBTB25, ZNF639 were amplified from Jurkat cDNA (RNeasy Plus kit; Qiagen) using the Q5 Hot Start DNA polymerase (NEB). TARBP1 was amplified from the plasmid pENTR223.1-hTARBP1, which was a gift from Falk Butter, using the Q5 Hot Start DNA polymerase (NEB) supplemented with the Q5 5X GC enhancer. The ZBTB2, ZBTB25, ZNF639 and TARBP1 inserts were then cloned into a retroviral pOZ-FH-C vector with a C-terminal FLAG and HA tag (FH) via restriction digest. Cloning of ZBTB2, ZBTB25 and ZNF639 was carried out with a ligation step using T4 ligase (Themo Fisher Scientific). Homologous recombination of the TARBP1 insert was employed to successfully clone the insert into the pOZ-FH-C vector (ClonExpress Ultra One Step Cloning Kit, Vazyme). Primers used are listed in **Supplementary Table 6**. Short-hairpin RNA (shRNA) were shortlisted from the GPP web portal (https://portals.broadinstitute.org/gpp/public/gene/search; BROAD institute) (see **Supplementary Table 6**) and were cloned into the pLKO.1-puro vector, following restriction enzyme digest and DNA ligation.

### Transfection and transduction

For retroviral and lentiviral production 300,000 HEK293T cells were seeded per well in a 6-well plate the evening prior to transfection. On the day of transfection 1 µg of the retroviral transfer vector was co-transfected with the 0.25µg of packaging vector pMD-MLV and 0.25 µg envelope vector pCMV-VSV-g into HEK293T cells using polyethylenimine (Polyscience) diluted in Opti-MEM. For lentivirus production, 0.5 µg of the transfer vector was co-transfected with 0.25 µg packaging vectors pMDLg/pRRE and pRSV-Rev, and 0.25 µg envelope vector pMD2.G into HEK293T cells using polyethylenimine (Polyscience) diluted in Opti-MEM. Media was changed 2 4h later to RPMI-1640 (Gibco). Viral supernatants were harvested with a 0.45 µm filter 72 h post-transfection. For viral transduction, viral supernatants were supplemented with 10 mM HEPES and 8 µg/mL polybrene (Sigma). The plate was spinoculated for 1.5 h at 2,500 rpm, 25-32 °C for efficient transduction. Media was changed 24 h later. For transduction of lentiviral shRNA, puromycin (0.75 µg/mL) was added 48 h post-transduction to select for transduced cells for another 72 h.

### qRT-PCR

Total RNA was extracted from cells using RNeasy Plus Mini Kit (Qiagen) as per manufacturer’s instruction. Superscript IV Reverse Transcriptase (Invitrogen) was used to generate cDNA from the extracted RNA. Transcript expression was measured by quantitative polymerase chain reaction (qPCR) with QuantiNova SYBR Green PCR Kit (Qiagen) in QuantStudio 3 or QuantStudio 5 Real-Time PCR System (Applied Biosystems) with a cycling condition of 95 °C for 2min, 95 °C for 5s and 60 °C for 30s for 40 cycles, followed by melting curve analysis. Transcript expression was assessed against the housekeeping gene TBP(TATA-binding protein) using the ΔΔCt method. Primers used are listed in **Supplementary Table 6**.

### Western blot

Cells were pelleted and washed once in 1x PBS before being resuspended in 2x Laemmli buffer (Sigma) and boiled for 5 min at 95°C. Alternatively, after a single wash of 1x PBS, the pellet was incubated with RIPA buffer (Sigma-Aldrich) supplemented with 1x cOmplete protease inhibitor (Roche) at 30 min on ice, followed by centrifugation for 30 min at maximum speed in 4ºC. Supernatant was transferred to a fresh tube and quantified with the Pierce BCA Protein Assay kit (Thermo Fisher Scientific). 30-40 µg of protein extract was diluted in 4x LDS sample buffer (Thermo Fisher Scientific) supplemented with 0.1M DTT and boiled for 10 min at 70ºC. Samples were separated on NuPAGE 4-12% Bis-Tris gels (Thermo Fisher Scientific) for 1 h at 170 V in 1x MOPS buffer (Thermo Fisher Scientific). Proteins were transferred onto a PVDF membrane (Bio-Rad) in 1x transfer buffer (25 mM Tris, 192 mM glycine, 0.02% SDS, 10% methanol) in a trans-blot SD semi-dry transfer cell (Bio-Rad) at 60-240mA for 1 h or in a power blotter semi-dry transfer station (Thermo Fisher Scientific) at 25V for 1 h. Proteins with a higher molecular weight such as TARBP1 required wet transfer using the X Cell II Blot Module (Thermo Fisher Scientific) at 35V for 2 h in 4ºC. Membranes were then blocked in blocking buffer (5% milk (Nacalai Tesque) in PBS-Tween), followed by primary and secondary antibody incubation. ECL substrate (Pierce ECL Western blotting substrate (Thermo Fisher Scientific) or Amersham ECL Prime Western blotting detection reagent (GE Healthcare) was applied to initiate the ECL reaction. The list of antibodies used are available in **Supplementary Table 11**.

### Azidohomoalanine (AHA)-Desthiobiotin Pulldown

Jurkat cells were washed once with 1x PBS before incubation with methionine-free RPMI (Gibco) supplemented with 10% dialyzed fetal bovine serum (PAN-Biotech) and 100 U/mL penicillin-streptomycin (Gibco) for 30 min prior to labelling. Jurkat cells were then incubated with 50 µM AHA (Thermo Fisher Scientific) in methionine-free RPMI for 4 h to label neo-synthesed proteins. Cells were harvested by centrifugation at 400 g for 5 min and washed thrice with 1x PBS. The pellet was lysed with 50 µL per 10^6^ cells of Lysis Buffer (50mM Tris-HCl pH 8.0, 1% SDS, 1X cOmplete protease inhibitor (Roche)) for 15 min on ice. DNA was then dispersed by mechanical shearing using the EpiShear Probe Sonicator (Active Motif) for 5-10 sec at 30% amplitude, followed by vortexing using a ThermoMixer (Eppendorf) at 1,500 rpm at RT for 5 min. The lysate was then centrifuged at 18,000 g for 5 min at 4ºC and the supernatant was transferred to a fresh tube and quantified with the Pierce BCA Protein Assay kit (Thermo Fisher Scientific). 200-600µg of protein lysate were then click-labelled with desthiobiotin alkyne (Vector Laboratories) using the Click-iT Protein Reaction Buffer Kit (Thermo Fisher Scientific) according to the manufacturer’s instructions. AHA-deshtiobiotin-tagged proteins are then precipitated via methanol-chloroform extraction and the pellet was dried from 15 min to overnight. The dried pellet was resuspended with 200-300µL 1xPBS containing 1% SDS and then sonicated using the EpiShear Probe Sonicator (Active Motif) for 10-20 sec at 30% amplitude followed by incubation in the ThermoMixer (Eppendorf) for 5 min at 1,500 rpm and 37ºC. 750 µg (WB) paramagnetic streptavidin beads (Dynabeads MyOne Streptavidin C1, Thermo Fisher Scientific) were pre-washed twice with PBB buffer containing 180 mM NaCl and 10 μg/μl BSA before resuspension with 150µL PBB buffer containing 420mM NaCl (thereafter the NaCl concentration remained the same for rest of the experiment). Beads were then incubated in 4ºC with rotation at 25 rpm for 2 h. Three PBB buffer washes were performed and bound proteins were eluted with 25 µL PBB containing 4mg/mL biotin (Sigma-Aldrich) at 37ºC for 30 min at 950 rpm. Finally, the protein elute was then boiled with 2x Laemmli buffer at 95ºC for 5 min.

## Data Availability

The mass spectrometry data have been deposited to the ProteomeXchange Consortium via the PRIDE partner repository^39^ with the dataset identifier PXD057962.

## Acknowledgments

We are grateful to all members of the Kappei lab for advice and discussions. Research in the Kappei lab was supported by the National Research Foundation Singapore and the Singapore Ministry of Education under its Research Centres of Excellence initiative and an NMRC Open Fund Young Individual Research Grant (NMRC/OFYIRG/055/2017). JC is supported by the Kay Kendall Leukaemia Fund. JC and MRM are funded through a CRUK Programme Foundation Award (DRCPFA\100012). MRM is supported by a Great Ormond Street Hospital Children’s Charity Professorship. We thank Diego Pitta de Araujo (NUHS-RSU) for the graphic design in Fig. 5D.

## Authorship contributions

NZA, CYG and DK conceived the study and designed experiments with input from all authors. NZA and CYG performed most experiments with help from BH and SHT. TS, JRC, and MM contributed critical reagents. NZA, CYG and DK analysed the data. NZA, CYG and DK wrote the manuscript with input from all authors.

## SUPPLEMENTARY FIGURES

**Figure S1.**
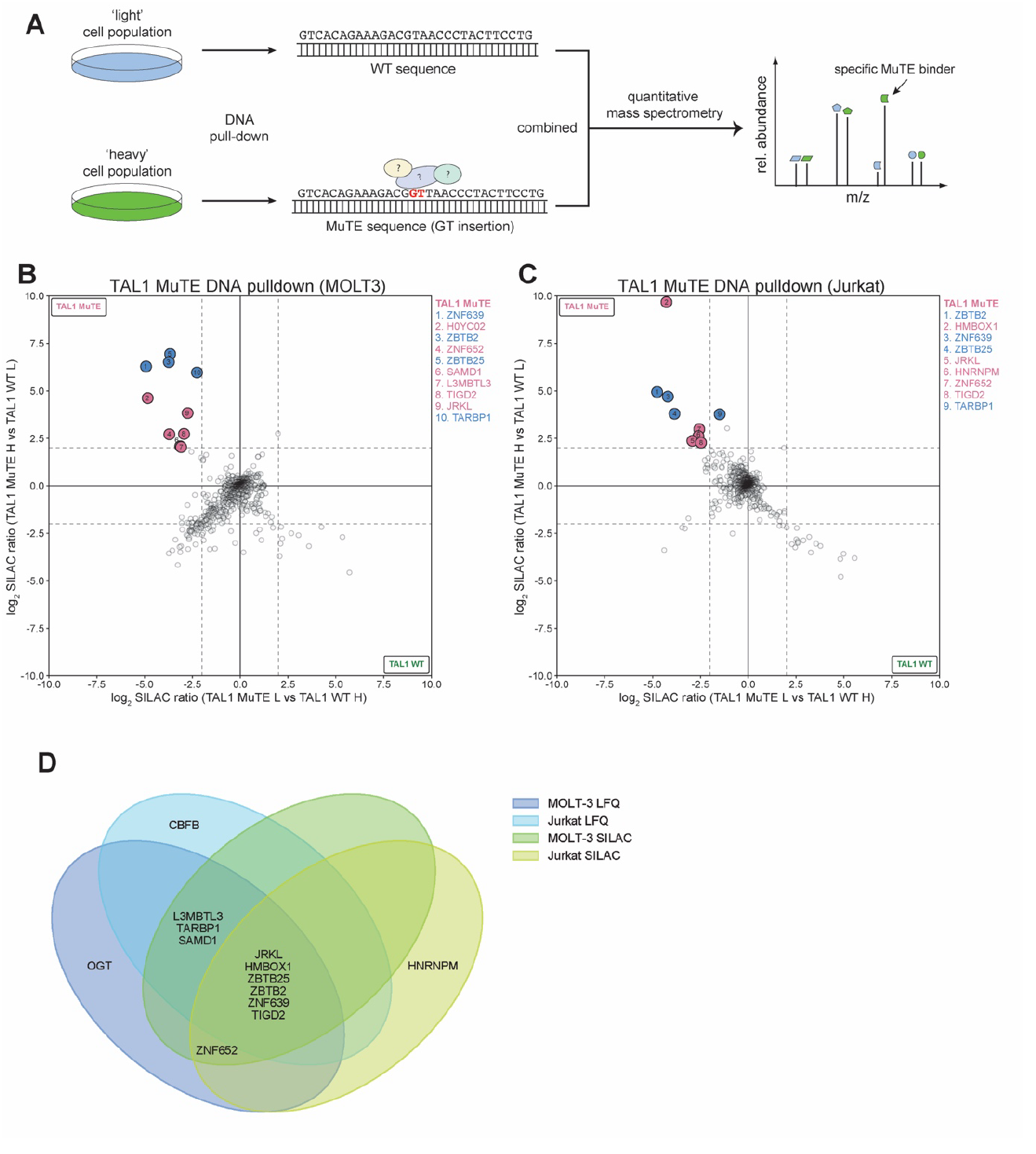
(A) *In vitro* reconstitution DNA pull-down combined with SILAC-based quantitative mass spectrometry. Differential SILAC-labelling is based on incubation of cells with either ‘heavy’ (^15^N- and ^13^C-labelled Lys and Arg) or ‘light’ (^14^N- and ^12^C-labelled Lys and Arg) medium. Heavy or light nuclear protein extracts are *in vitro* reconstituted with either WT or MuTE probes. The two pull-down fractions are combined and analysed in the mass spectrometer. Relative enrichment of heavy and light peptides (“SILAC pairs”) quantitatively measures the enrichment of specific binding proteins to either of the two sequences. (B) Scatterplot of DNA pull-downs with SILAC-labelled nuclear protein extracts from MOLT-3 and (C) Jurkat cells. Heavy-labelled SILAC extracts were applied to the MuTE oligonucleotides and light-labelled extracts to the WT oligonucleotides (y-axis) or vice versa (x-axis) and log_2_ SILAC ratios are displayed. Specific MuTE-binders are found in the upper left quadrant as numbered (cut-off ≥4-fold enrichment). (D) Venn diagram showing overlap of MuTE-specific proteins identified across four DNA pull-down-MS datasets.

**Figure S2.**
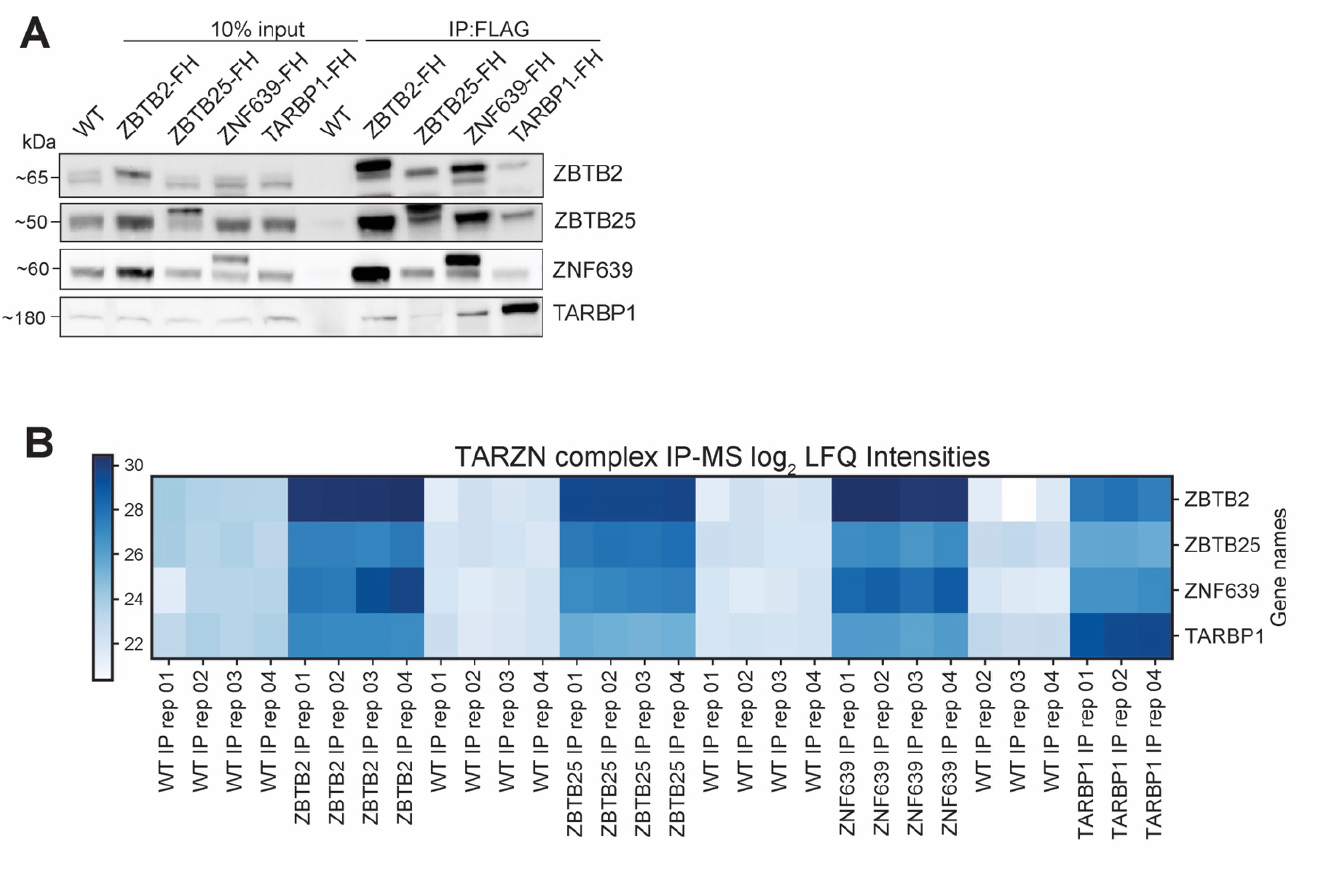
(A) Immunoblot validating IP-MS in Fig. 3B-D. Blots shown are representative results from at least three independent experiments. Antibodies against each respective protein (ZBTB2, ZBTB25, ZNF639 and TARBP1) were used, where a double band pattern is visible for the overexpression constructs. Specifically, the lower molecular weight band represents the endogenously expressed protein and the shift of the higher molecular weight band represents the overexpressed protein with the additional FLAG-tag. (B) Heatmap of TARZN complex IP-MS log_2_ LFQ intensities.

**Figure S3.**
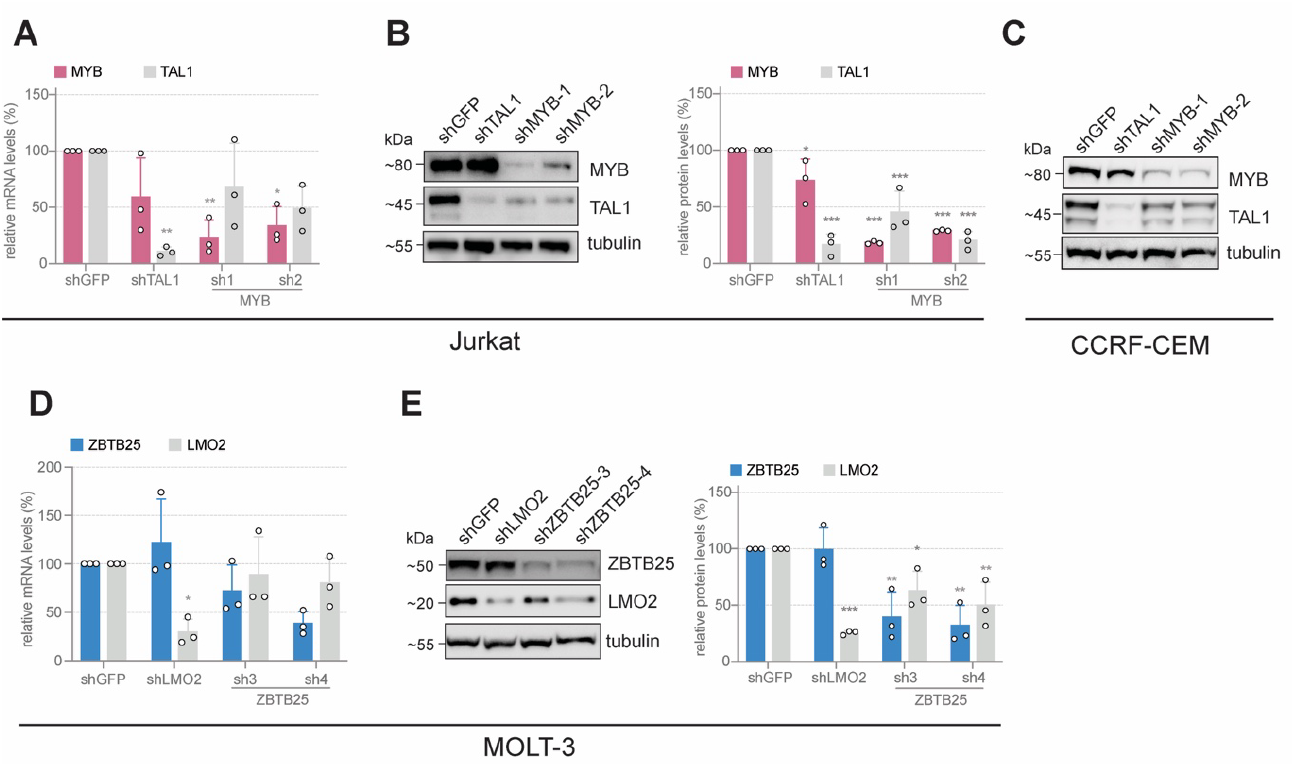
(A) qPCR data measuring *MYB* knockdown and its effect on *TAL1* expression at the mRNA level in Jurkat cells. Cells were transduced with lentivirus expressing shRNAs of the target proteins, and puromycin selection (0.75 µg/ml) was initiated 48 h post-transduction for a further 72 h. (B) Immunoblots showing shRNA-mediated knockdown of MYB in Jurkat cells and its effect on TAL1 protein expression. Cells were transduced with lentivirus expressing shRNAs of the target proteins, and puromycin selection (0.75 µg/ml) was initiated 48 h post-transduction for a further 72 h. Whole cell lysates were used as the input for immunoblotting. Right panel shows quantification of the immunoblots from Jurkat cells performed using ImageJ. (C) Immunoblots showing shRNA-mediated knockdown of MYB in MuTE-negative CCRF-CEM cells and <u>its</u> effect on TAL1 protein expression. Experiment was performed as described for Jurkat cells in (A). (D) qPCR data measuring *ZBTB25* knockdown and its effect on *LMO2* expression at the mRNA level in MOLT-3 cells. Cells were transduced with lentivirus expressing shRNAs of the target proteins, and puromycin selection (0.75 µg/ml) was initiated 48 h post-transduction for a further 72 h. (E) Immunoblots showing shRNA-mediated knockdown of ZBTB25 in MOLT3 cells and their effect on LMO2 protein expression. Experiment was performed as described for Jurkat cells in (A). All data shown depicts the mean±SD of three biological replicates. Asterisks represent statistical significance (^*^ p≤0.05, ^**^ p≤0.01, ^***^ p≤0.001) calculated by one-way ANOVA followed by Dunnett’s multiple comparison tests.

